# Style Transfer Using Generative Adversarial Networks for Multi-Site MRI Harmonization

**DOI:** 10.1101/2021.03.17.435892

**Authors:** Mengting Liu, Piyush Maiti, Sophia Thomopoulos, Alyssa Zhu, Yaqiong Chai, Hosung Kim, Neda Jahanshad

## Abstract

Large data initiatives and high-powered brain imaging analyses require the pooling of MR images acquired across multiple scanners, often using different protocols. Prospective cross-site harmonization often involves the use of a phantom or traveling subjects. However, as more datasets are becoming publicly available, there is a growing need for retrospective harmonization, pooling data from sites not originally coordinated together. Several retrospective harmonization techniques have shown promise in removing cross-site image variation. However, most unsupervised methods cannot distinguish between image-acquisition based variability and cross-site population variability, so they require that datasets contain subjects or patient groups with similar clinical or demographic information. To overcome this limitation, we consider cross-site MRI image harmonization as a style transfer problem rather than a domain transfer problem. Using a fully unsupervised deep-learning framework based on a generative adversarial network (GAN), we show that MR images can be harmonized by inserting the style information encoded from a reference image directly, without knowing their site/scanner labels *a priori*. We trained our model using data from five large-scale multi-site datasets with varied demographics. Results demonstrated that our style-encoding model can harmonize MR images, and match intensity profiles, successfully, without relying on traveling subjects. This model also avoids the need to control for clinical, diagnostic, or demographic information. Moreover, we further demonstrated that if we included diverse enough images into the training set, our method successfully harmonized MR images collected from unseen scanners and protocols, suggesting a promising novel tool for ongoing collaborative studies.

## 1 Introduction

Large-scale multi-site neuroimaging studies allow for high statistical power and are essential for ensuring reliable and robust results. Many multi-site initiatives prospectively aim to harmonize image acquisition protocols to minimize scanner-induced variability due to factors such as magnetic field strength, coil channels, gradient directions, manufacturer, and image resolution. However, in many cases, the need for retrospective harmonization is often inevitable. Decade-long running studies, such as ADNI undergo scanner upgrades, and many studies of small effects or effects across wide demographic ranges, require retrospective pooling of data.

Several retrospective image harmonization techniques have shown promise in being able to remove cross-site variance from different studies for pooled analyses. Most harmonization methods fall into two broad categories: 1) harmonization of image-derived features using statistical properties of the distribution, for example ComBat (Fortin et al., 2017); 2) harmonization of the task output of the MR images, i.e. segmentation, classification, and prediction. This second category is largely composed of deep learning-based approaches, namely domain adaptation techniques. In particular, domain transfer learning and domain adversarial learning have been successfully applied for MRI harmonization (Dinsdale et al., 2020).

However, if a wide range of tasks were to be performed on images, harmonization would need to be performed separately for each task. Furthermore, there are some applications, like cortical surface construction (as opposed to segmentation), that cannot be directly embedded into a deep learning framework (Dong et al., 2020). In these scenarios, image translation or a direct image-to-image harmonization for MRIs is needed. Several image translation-based harmonization methods have been proposed. Supervised methods typically require traveling subjects and must be planned prospectively (Dewey, et al., 2019). Unsupervised methods, such as variational auto-encoders (Moyer et al., 2020) or CycleGAN (Zhu et al., 2017), MR images are often separated into well-defined domains in terms of scanners or sites. These methods are prone to overcorrection if data collected at each site are not acquired with different scanning parameters, but are also confounded by differences in population of individuals scanned, such as clinical diagnoses, age ranges, or even race and ethnic groups; the correction for site could then also remove critical biological differences. In these situations, the demographic and clinical conditions need to be strictly controlled and matched. Furthermore, even with the data collected from the same scanners/sites, the images collected may also show slight variance (Tong et al., 2020). These intra-site differences make the harmonization less accurate due to the lack of diversity of these domain-based methods.

Recently, deep learning methods have successfully completed diverse image translations by disentangling the image into ‘content’ and ‘style’ spaces (Huang et al., 2018), where contents represent low level information in images like contours and orientations, and styles can be considered high level information such as colors and textures. Images within the same domain share the same content space but may show different styles. In MR images, we can consider the biologically defined anatomical information as the content, and the non-biological information such as intensity variation, SNR, and contrasts as styles. Dewey, et al., (2020) have used this breakdown to show promising results for MRI harmonization. However, in (Dewey et al., 2020) paired image modalities from the same subjects were needed to supervise the extraction of the content information.

Here, we consider image harmonization as a pure style transfer problem rather than a domain transfer problem. We think anatomical patterns (or contents) from MR images collected from different sites share the same latent space, and it is not necessary to separate them into different “domains’’. These style-irrelevant patterns can be learned using an unsupervised cycle consistency GAN model, and thus any paired modalities are not needed from the same subjects. Due to scanner shifts, and software upgrades, the styles for all the images, even collected from the same site may be different from each other. Furthermore, inspired by (Choi et al., 2020), we proposed that the style information needed for harmonization can be encoded from a single reference MR image directly, instead of using predefined labels.

## 2 Method

### 2.1 The Architecture of Style-Encoding GAN

Let *X* be the set of MR images. Given an image *x* ∈ *X*, our goal is to train a single generator *G* that can generate diverse images that correspond to the image *x* with a style code *s*, where *s* is associated with the style (non-biological) patterns from another image. The style code *s* is generated by a mapping network *M* from sampling a given latent vector *z* (*s* = *M*(*z*)). The reason for using *s* instead of *z* has been provided by past studies (Karras et al., 2019). Then, the generator *G* translates an input image *x* into an output image *G*(*x*, *s*) that reflects the style of *s*. To validate that the style code *s* is successfully injected into the output image *G*(*x*, *s*), another style encoding network *E* was designed to encode the style of *s* from images. That is, given an image *x*, the encoder *E* extracts the style code *s* = *E*(*x*) of *x*. *E* can produce diverse style codes using different images. This allows *G* to synthesize an output image reflecting the style *s* from different reference images of *X*. The goal of the network is to train *E* so that *E*(*G*(*x, s*)) = *s*, meaning that if an image was generated based on style code *s*, then *s* can also be encoded when this image was input into the style encoder *E*. Adaptive instance normalization (AdaIN) (Huang et al., 2017) was used to inject *s* into *G*. Finally, the discriminator *D* learns a binary classification determining whether an image *x* is a real image or a fake image *G*(*x*, *s*) produced by *G*. Borrowed by (Choi et al., 2018), our model includes only one generator, one discriminator, and one style encoder.

### 2.2 Network Training

Given an image *x* ∈ *X*, we train our framework using the following objectives.

#### Adversarial loss

During training, we sample a latent code *z* ∈ *Z* randomly, and the mapping network *M* learns to generate a target style code *s* = *M* (*z*). The generator *G* takes an image *x* and *s* as inputs and learns to generate an output image *G*(*x*, *s*) that is indistinguishable by the discriminator *D* from real images via an adversarial loss:

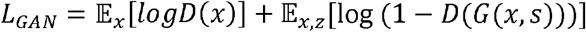

#### Cycle-Consistency Loss

To guarantee that generated images are meaningful to the original images and properly preserving the style-irrelevant characteristics (e.g. anatomical patterns) of input *x*, an additional cycle consistency loss (Zhao et al., 2019) is defined as the difference between original and reconstructed images:

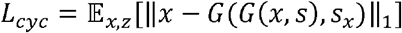

where *s_x_* = *E*(*x*) is the estimated style code of the input image *x*. By encouraging the generator *G* to reconstruct the input image *x* with the estimated style code *s_x_, G* learns to preserve the original characteristics of *x* while changing its style faithfully.

#### Style reconstruction loss

In order to enforce the generator *G* to use the style code while generating the image *G*(*x, s*), we incorporate a style reconstruction loss:

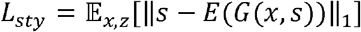

Our learned encoder E allows G to transform an input image *x*, to reflect the style of a reference image.

#### Style diversification loss

To further enable the generator *G* to produce diverse images, we explicitly regularize *G* with the diversity sensitive loss (Wang et al., 2018):

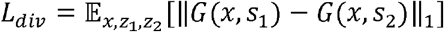

where the target style codes *s*_1_ and *s*_2_ are produced by *M* conditioned on two random latent codes *z*_1_ and *z*_2_(i.e. *s_i_* = *M*(*z_i_*) for *i* ∈ {1, 2}). Maximizing the regularization term forces *G* to explore the image space and discover meaningful style features to generate diverse images.

Put together, our full objective function can be summarized as follows:

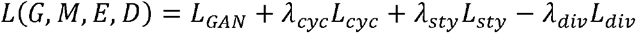

Where *λ_cyc_, λ_sty_* and *λ_div_* are hyperparameters for each term.

### 2.3 Experimental Setup

We obtained T1-weighted brain MR images from five datasets: UK Biobank (UKBB), Parkinson’s Progression Markers Initiative (PPMI), and Alzheimer Disease Neuroimaging Initiative (ADNI), Adolescent Brain Cognitive Development (ABCD) and International Consortium for Brain Mapping (ICBM). Scans used in this study were collected from disease-free participants (UKBB: n=200, age range 45-55 years old; PPMI: n=76 age range 67-70 years old; ADNI: n=42, age range 67-70 years old; ABCD: n=200, age range 9-13 years old; and ICBM: n=200, age range 19-54 years old; Fig. 2), among which 90% were used as training/validation sets and 10% testing sets. We deliberately kept some cohorts overlapping completely in age, while others were non-overlapping. To show that our model generalizes to unseen images, we also applied the trained harmonization model to one travelling subject collected in (Tong et al., 2020), who was scanned 12 times at 10 different sites within 13 months. This traveling subject was also used to validate the harmonization quantitatively. All image acquisition information for these public resources can be found elsewhere, but briefly they vary in terms of scanner manufacturer, field strength, voxel size, and more, often within the same study.

**Fig. 1.**
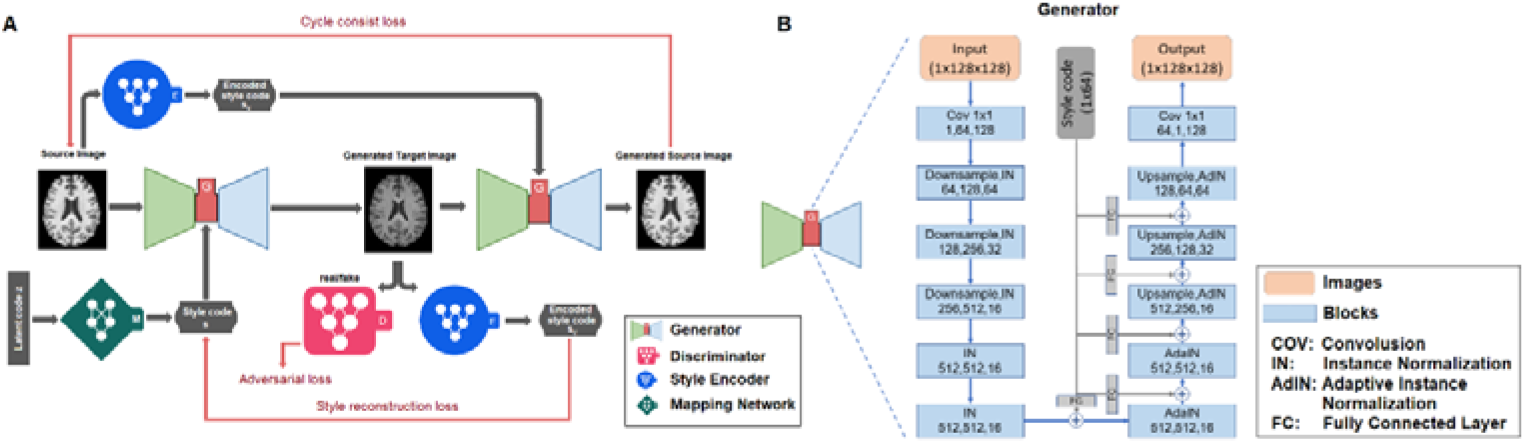
A) The architecture of the style-encoding GAN. Generators learns to generate image by inputting a source image and a style code. The learning process is driven by cycle consist loss, adversarial loss, style reconstruction loss and style diversification loss (not shown in this figure) B) The detailed architecture of the generator in network. In each of the block, the three number means number of input channels, number of output channels and the image size.

**Fig. 2.**
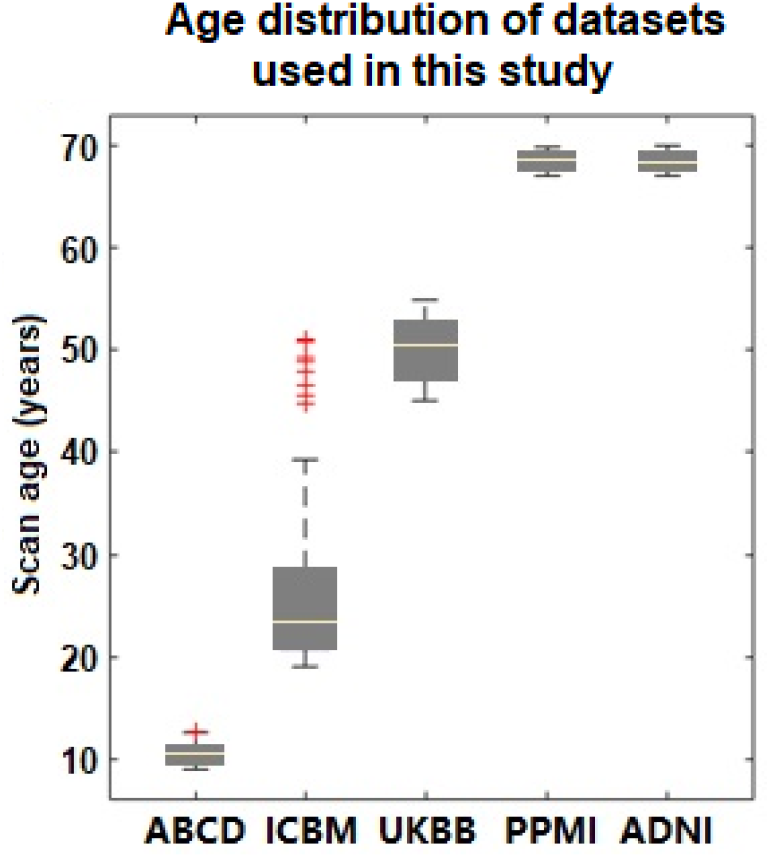
Age distribution of the five datasets used in this study. Very limited age overlap were applied.

All the images were skull-stripped, nonuniformity corrected and registered to the MNI template using a 9 dof linear registration. MR images from the UKBB and PPMI datasets were further segmented into white matter (WM), gray matter (GM) and cerebrospinal fluid (CSF) using a statistical partial volume model (Tohka et al., 2004). All the images were resized to 128×128×128 voxels. To pilot this work, we used 2D slices as input, and we selected 50 axial slices in the middle of each MRI volume as input.

## 3 Results

### 3.1 Validation of the Removal of Cross-sites Variances

#### Histogram Comparison

Fig. 3 illustrates the harmonized images among the 5 datasets according to nine randomly selected reference images from across the datasets. Harmonized images are noticeably more similar in contrast and intensity to those of the reference images, and their anatomical structures were well-maintained. To quantitatively compare the intensity histogram of the harmonized images and reference images, we random select one participant from ADNI, and one participant from PPMI. Participants were matched for age and sex to ensure approximately similar volumes of each tissue type. We made a slice-matched comparison between the histograms of all 50 slices in MR images from the two participants, and the histogram of the image after the ADNI scan after it was translated to the PPMI domain using Jensen-Shannon divergence (JS). The JS between ADNI and translated ADNI→PPMI image () is significantly higher than JS between PPMI and the ADNI→PPMI translated image (; p<0.0001), suggesting the histogram of translated image has an intensity profile more similar to PPMI than ADNI, from which it came.

**Fig. 3.**
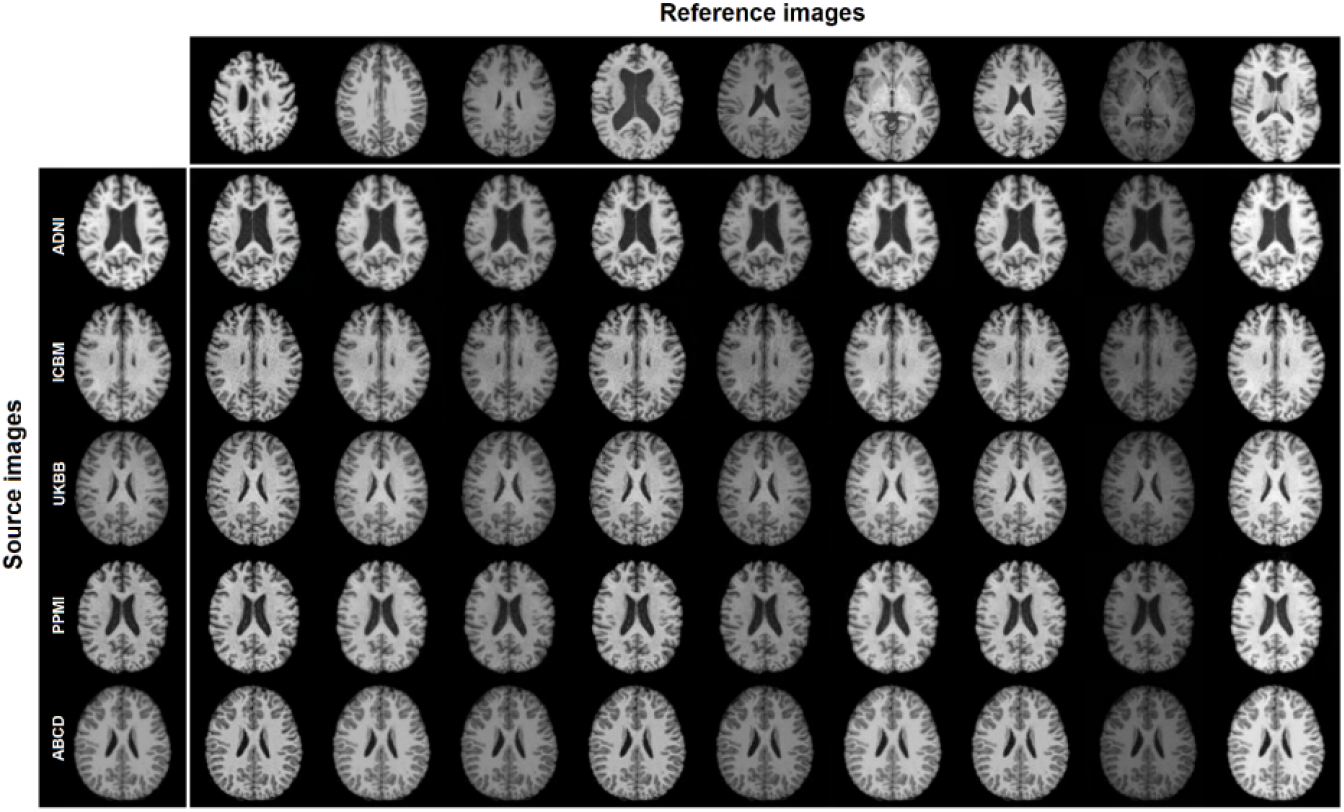
The style-encoding GAN can harmonize images based on only one single reference image.

#### Segmentation Comparison

Brain segmentation is usually sensitive to image contrast variations, which may result in systematic over- and under-segmentation of brain tissues in a way that is inconsistent across acquisition protocols. Here we further validate the removal of cross-site variances by testing a segmentation model for images before and after the harmonization. In this example, MR slices from UKBB dataset (10% testing set as described in the harmonization section) were used to train a deep-learning based U-net to segment the brain into WM, GM, and CSF. The auto-segmentation outputs were used as the ground truth. The trained model was then applied to slices in the PPMI test dataset (testing set in the harmonization section) for segmentation before and after the harmonization using a reference image from UKBB dataset. The Dice score for WM (before vs. after harmonization: 0.93±0.02 vs. 0.95±0.01), GM (0.80±0.06 vs. 0.86±0.02) and CSF (0.51±0.16 vs. 0.64±0.14) were all significantly higher for images after harmonization (all p < 0.0001).

### 3.2 Validation of the Preserve of Brain Anatomical Information

Ideally, MR image harmonization methods not only remove the cross-sites variances, but also rigorously maintain the anatomical information within subjects. To test whether the anatomical information in MR images were preserved after harmonization, we compared the intra-subject similarity and inter-subject differences before and after the harmonization. In this illustration, to guarantee the similarity between slices, we selected only the middle slices from 10 subjects randomly from ADNI data as source images, and 100 images randomly selected from other datasets as reference images.

#### Preserve of intra-subject anatomical patterns

We compared the similarity between images before and after harmonization to different scans using intensity correlation (r) and the structural similarity index measure (SSIM) as metrics. The average intensity correlation between images before and after harmonization is 0.984 and the SSIM is 0.612. Both values were significantly higher than the average value between pairs of images before harmonization (r = 0.901; SSIM = 0.412), indicating the intra-subject anatomical information was preserved after the harmonization compared to inter-subject variances.

#### Preserve of cross-subject differences

To quantify inter-subject differences, we used the intensity differences to compute Euclidean distances (Zhao et al., 2019) between any two scans, forming a distance matrix, denoted as 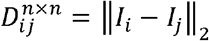, where n = 10 is the number of scans, and *I* the whole-image voxel intensity vectors for scans *i* and *j*. The goal was to estimate how the distances were preserved relative to each other before and after harmonization. We computed the correlation r between the two distance-matrices before and after harmonization. Our model achieves an average Cor of 0.963 (range: [0.943, 0.991]) between the distance-matrices before harmonization and the 100 distance-matrices after harmonization, indicating the intersubject difference was reliably preserved after harmonization.

### 3.3 Hyperparameter Selection

If the groups of images to be harmonized are confounded by demographic or clinical/pathological differences, such biological differences could also be inadvertently learned during the harmonization. To avoid this, we tuned the in our model to preserve the style-irrelevant characteristics. Fig. 4 shows an example of the image harmonized using the model with different values. In this example, the source image is from ADNI and the reference image is from ABCD. There is approximately a 58-year age difference between these subjects. If =0, meaning none of the style-irrelevant characteristics are needed, the model learns everything from the reference image, generating an image completely identical to the reference. If, then the model learns style from the reference image but also some biological patterns, such as smaller lateral ventricles and thicker gray matter cortices. If, then the model learns only the style information from the reference and rigorously maintains the style-irrelevant characteristics (ventricles etc.) from the source images.

**Fig. 4.**
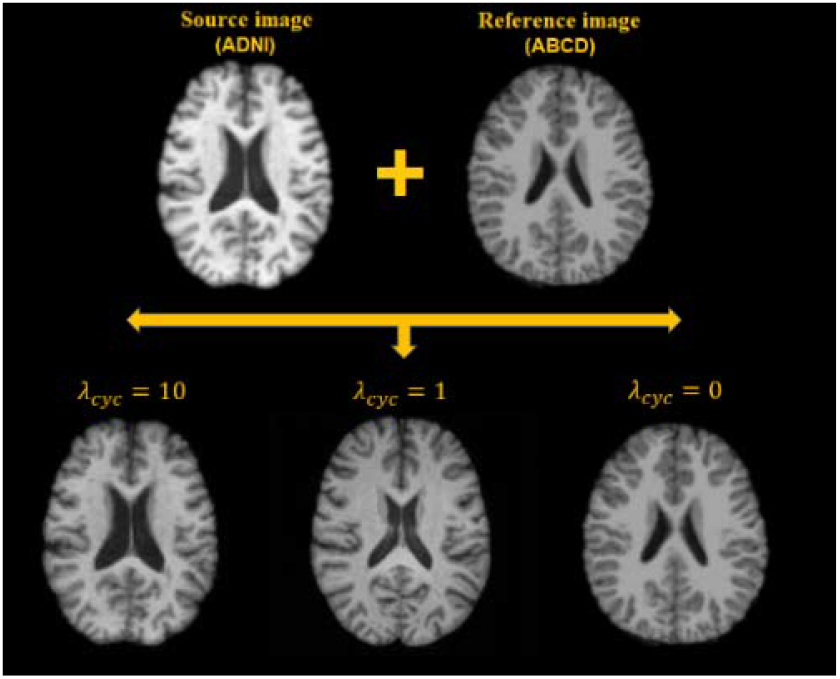
Small cycle consistency loss coefficients bias the generated images towards the reference images, while larger cycle consistency coefficients rigorously maintain the structure of the source images.

### 3.4 Harmonization for MR Images from unseen dataset

By design, the model should learn style-invariant features, encouraging better generalization to unseen samples. Fig. 5 shows that our model successfully captures styles of the unseen/traveling subject and renders these styles correctly to the source images. Furthermore, we harmonize all the 12 scans from the traveling subject using one randomly selected scan (among 12 scans) as the reference. We then compared the slice-matched similarity between all paired scans using SSIM and peak signal to noise ratio (PSNR), and find a significant improvement in similarity using SSIM (M = 0.752 before harmonization vs. 0.822 for harmonized images, two-way t-test: p = 0.031). PSNR also showed an improved similarity but was not statistically significant (23.1 before harmonization and 24.1 after harmonization).

**Fig. 5.**
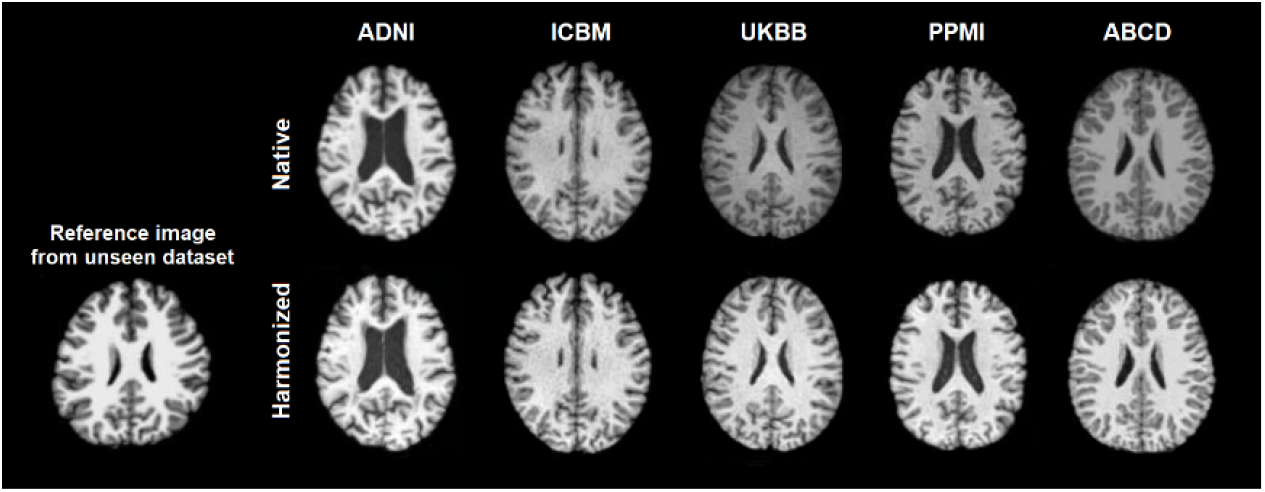
The trained style-encoding GAN successfully captures styles of reference images from novel acquisition protocols and renders these styles correctly to the source images.

## 4 Discussion and Conclusion

We have shown in this study that the style-encoding GAN paves a way for multi-site MR images to be synthesized from a single input image from arbitrary sites, taking the style code encoded from an arbitrary reference image directly as extra information, without knowing the acquisition protocols *a priori*. Our model does not rely on traveling subjects or any paired modalities of images from the same subjects. Furthermore, because we consider the cross-site harmonization as a style transfer problem rather than a domain transfer problem, MR images from multiple sites need not be categorized into different domains. Thus, the demographic and pathological conditions do not need to be matched for the harmonization. We showed our model generalizes well even to unseen samples, likely due to the diverse data included in the training set. This is particularly important for individual studies with very small sample sizes using acquisition protocols that are seldomly used by other datasets.

One of the limitations in applying this model is how to select the best reference image if large-scale harmonization was applied. Also, currently the method only works on 2-D slices, future work will focus on how to harmonize images on 3-D spaces using the current approach.

## Acknowledgements

This work was supported in part by: R01AG059874, RF1AG057892, U01AG068057, and P41EB015922. BrightFocus Research Grant award (A2019052S). This research has been conducted using the UK Biobank Resource under Application Number ‘11559’. Data used in the preparation of this article were also obtained from the Parkinson’s Progression Markers Initiative (PPMI) database (www.ppmi-info.org/data), the Alzheimer’s Disease Neuroimaging Initiative (ADNI) database (adni.loni.usc.edu), and the Adolescent Brain Cognitive Development (ABCD) Study (https://abcdstudy.org), held in the NIMH Data Archive (NDA).

